# An effective response to respiratory inhibition by a *Pseudomonas aeruginosa* excreted quinoline promotes *Staphylococcus aureus* fitness and survival in co-culture

**DOI:** 10.1101/2025.03.12.642861

**Authors:** Franklin Roman-Rodriguez, Jisun Kim, Dane Parker, Jeffrey M. Boyd

**Affiliations:** Department of Biochemistry and Microbiology, Rutgers, The State University of New Jersey, New Brunswick, NJ 08901, USA; Department of Pathology, Immunology and Laboratory Medicine, Center for Immunity and Inflammation, Rutgers New Jersey Medical School, Newark, NJ 07103, USA

**Keywords:** SrrAB, Rex, respiration, HQNO, *Pseudomonas aeruginosa*, *Staphylococcus aureus*

## Abstract

*Pseudomonas aeruginosa* and *Staphylococcus aureus* are primary bacterial pathogens isolated from the airways of cystic fibrosis patients. *P. aeruginosa* produces secondary metabolites that negatively impact the fitness of *S. aureus,* allowing *P. aeruginosa* to become the most prominent bacterium when the species are co-cultured. Some of these metabolites inhibit *S. aureus* respiration. SrrAB is a staphylococcal two-component regulatory system (TCRS) that responds to alterations in respiratory status and helps *S. aureus* transition between fermentative and respiratory metabolisms. We used *P. aeruginosa* mutant strains and chemical genetics to demonstrate that *P. aeruginosa* secondary metabolites, HQNO in particular, inhibit *S. aureus* respiration, resulting in modified SrrAB stimulation. Metabolomic analyses found that the ratio of NAD^+^ to NADH increased upon prolonged culture with HQNO. Consistent with this, the activity of the Rex transcriptional regulator, which senses and responds to alterations in the NAD^+^ / NADH ratio, had altered activity upon HQNO treatment. The presence of SrrAB increased fitness when cultured with HQNO and increased survival when challenged with *P. aeruginosa. S. aureus* strains with a decreased ability to maintain redox homeostasis via fermentation had decreased fitness when challenged with HQNO and decreased survival when challenged with *P. aeruginosa*. These findings led to a model wherein *P. aeruginosa* secreted HQNO inhibits *S. aureus* respiration, stimulating SrrAB, which promotes fitness and survival by increasing carbon flux through fermentative pathways to maintain redox homeostasis.

**Importance:** Cystic fibrosis (CF) is a hereditary respiratory disease that predisposes patients to bacterial infections, primarily caused by *Staphylococcus aureus* and *Pseudomonas aeruginosa*. *P. aeruginosa* excreted secondary metabolites decrease *S. aureus* fitness during co-infection, ultimately eliminating it. The genetic mechanisms that *S. aureus* uses to detect and respond to these metabolites are unknown. The *S. aureus* SrrAB two-component regulatory system senses flux through respiratory pathways and increases transcription of genes utilized for adaption to low-respiration environments. This study demonstrates that SrrAB responds to the *P. aeruginosa*-produced respiratory toxin HQNO and responds by increasing fermentation increasing competition. This study describes interactions between these two bacterial pathogens, which could be exploited to decrease pathogen burden in individuals living with cystic fibrosis.

## Introduction

Cystic fibrosis patients suffer from increased lung and airway mucosa because of dysfunctional cystic fibrosis transmembrane conductance regulator (CFTR) (1). CFTR helps transport chloride through cell membranes, but when its function is disrupted, less chloride is exported from epithelial cells, leading to thicker mucus in the lungs, which is difficult to clear, serving as a less hostile environment for bacteria (1). Cystic fibrosis patients often have polymicrobial lung infections, with *S. aureus* and *P. aeruginosa* being the two most abundant bacteria (2). Methicillin-resistant *S. aureus* (MRSA) is found in around 20% of individuals with cystic fibrosis between the ages of 10-20, and *P. aeruginosa* is found in around 17% of patients, and 3.5% of these carry multi-antibiotic-resistant strains (2). *S. aureus* is most prevalent in children and teenagers, but as patients age, *P. aeruginosa* becomes the most prevalent, coinciding with a worsening of symptoms and lower lung activity (2–8). *P. aeruginosa* secretes various compounds that cause *S. aureus* cell lysis, inhibition of respiration, and iron starvation, allowing it to outcompete *S. aureus* (9–15). Co-culture of *P. aeruginosa* with *S. aureus* can select for *S. aureus* small colony variants (SCV) (16). SCVs are respiration deficient and rely on fermentation for energy generation. A fermentative lifestyle increases lactate release, which is conversely used as a primary carbon source by *P. aeruginosa* during co-infections (11, 16, 17).

The secretion of multiple *P. aeruginosa* secondary metabolites is regulated by quorum sensing systems. There is a total of four described quorum sensing (QS) systems in *P. aeruginosa*, with the *Pseudomonas* quinolone system (PQS) being central to the production of various virulence factors (18). The *pqsABCDE* operon is one of the three main gene clusters of the PQS system, with the transcriptional regulator PqsR acting as the receptor and regulating transcription (Fig. 1) (18, 19). The PQS QS system is responsible for regulating the production of several virulence factors like 2-Heptyl-4-Quinolone N-Oxide (HQNO), pyocyanin, elastase, and rhamnolipids (9–14, 20). While these molecules are known to interfere with *S. aureus* physiology, the exact mechanisms of action have not been elucidated.

**Figure 1.**
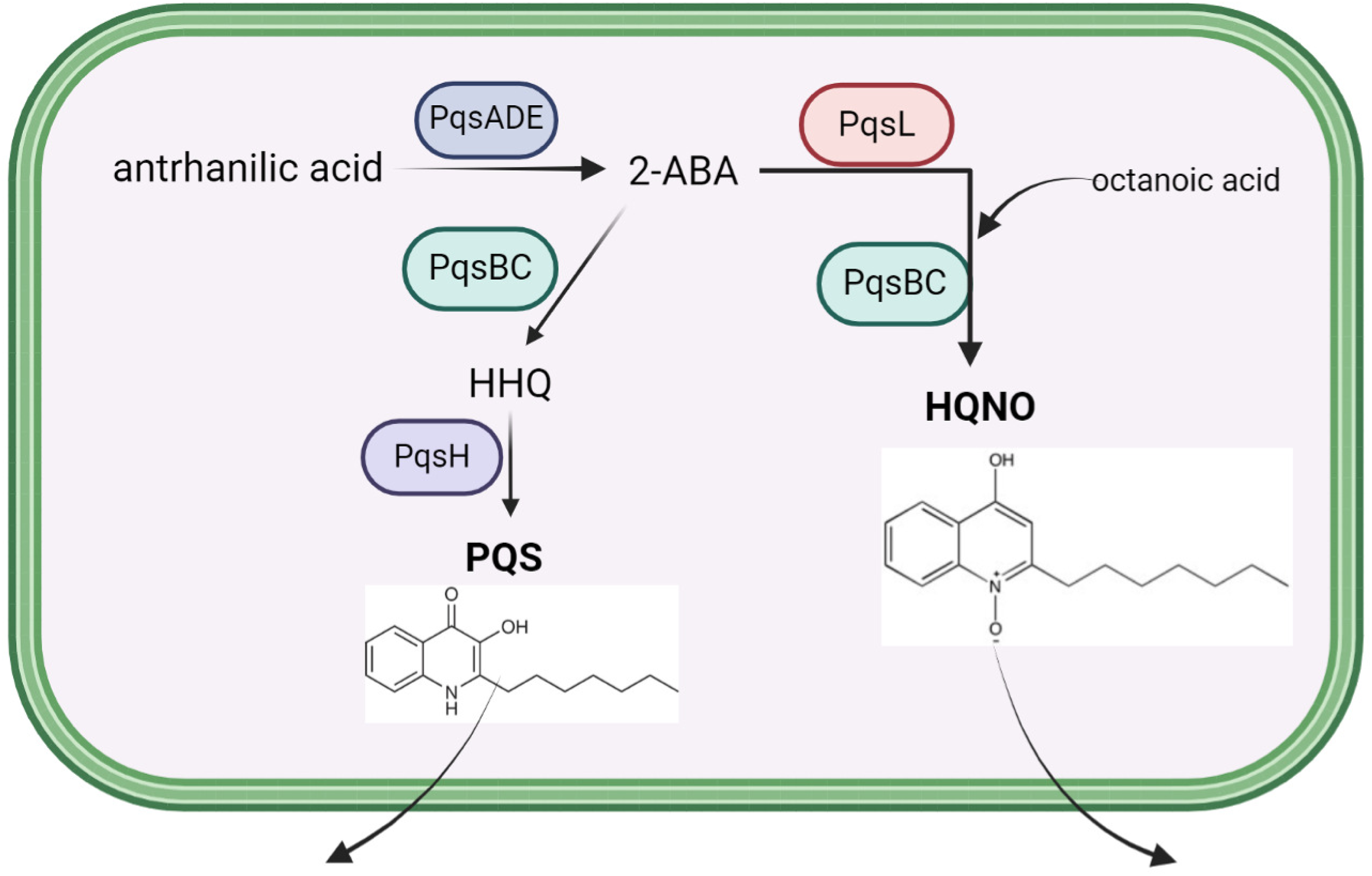
PQS and HQNO synthesis in *P. aeruginosa*. The PQS system of *P. aeruginosa* produces the secondary metabolites PQS and HQNO from anthranilic acid. The names of the enzymes responsible for the transformations are also displayed.

The staphylococcal respiratory response regulator (SrrAB) is a two-component regulatory system (TCRS) that controls the transcription of genes utilized for respiratory and fermentative metabolisms (21–23). SrrB is a membrane-spanning histidine kinase/phosphatase (HK), and SrrA is a cytosolic response regulator (RR). The HK responds to a stimulus by altering the phosphorylation status of the RR. The RR has DNA binding activity, and the affinity for DNA is altered by phosphorylation. The exact environmental and metabolic stimuli of the SrrAB system are unknown, but it is believed to sense the redox status of the menaquinone pool. SrrAB has increased activity during conditions where reduced menaquinone accumulates, and transcription of SrrAB-regulated genes is diminished in a menaquinone auxotroph (21). Upon decreased respiration, SrrAB increases transcription of the genes coding the Cyd and Qox terminal oxidases, as well as genes coding the anaerobic ribonucleotide reductase (Nrd), pyruvate formate lyase (Pfl), and lactate dehydrogenases (Ldh) (24, 25).

We tested the hypothesis that the S. aureus SrrAB TCRS responds to respiratory inhibition caused by *P. aeruginosa* secreted secondary metabolites. Data presented demonstrate that SrrAB senses HQNO-dependent respiratory inhibition, and the presence of SrrAB increases survival when challenged with *P. aeruginosa*. SrrAB-regulated genes that code for enzymes used for fermentation had decreased fitness upon HQNO challenge and decreased survival when co-cultured with *P. aeruginosa*. Furthermore, we demonstrate that HQNO treatment of *S. aureus* increases the ratio of NAD^+^ to NADH, resulting in altered activity of the Rex transcriptional regulator. These findings provide a framework for further investigations into the physiological changes that *S. aureus* undergoes when challenged with *P. aeruginosa*.

## Results

### *P. aeruginosa* secreted secondary metabolites stimulate SrrAB and alter *srrAB* transcriptional activity independent of SrrAB

We tested the hypothesis that *P. aeruginosa* secreted secondary metabolites stimulate SrrAB. The gene coding the iron-sulfur cluster requiring ribonucleotide reductase (*nrdD*) is transcriptionally regulated by SrrAB. It is one of the most upregulated genes when *S. aureus* transitions from respiratory to fermentative growth (24). Cell-free spent culture medium isolated after growth of wild type *P. aeruginosa* (PA14) was added to liquid cultures of *S. aureus* USA300_LAC (wild type or WT) or the isogenic Δ*srrAB* mutant. Both strains contained a *nrdD* transcriptional reporter. There was a significant increase in *nrdD* transcriptional activity upon co-culture with the PA14 spent medium (Fig 2A). Transcriptional activity was below the detectable limit in the Δ*srrAB* mutant. The data suggest that SrrAB is required to transcribe *nrdD* and that a *P. aeruginosa*-secreted metabolite is stimulating SrrAB.

**Figure 2.**
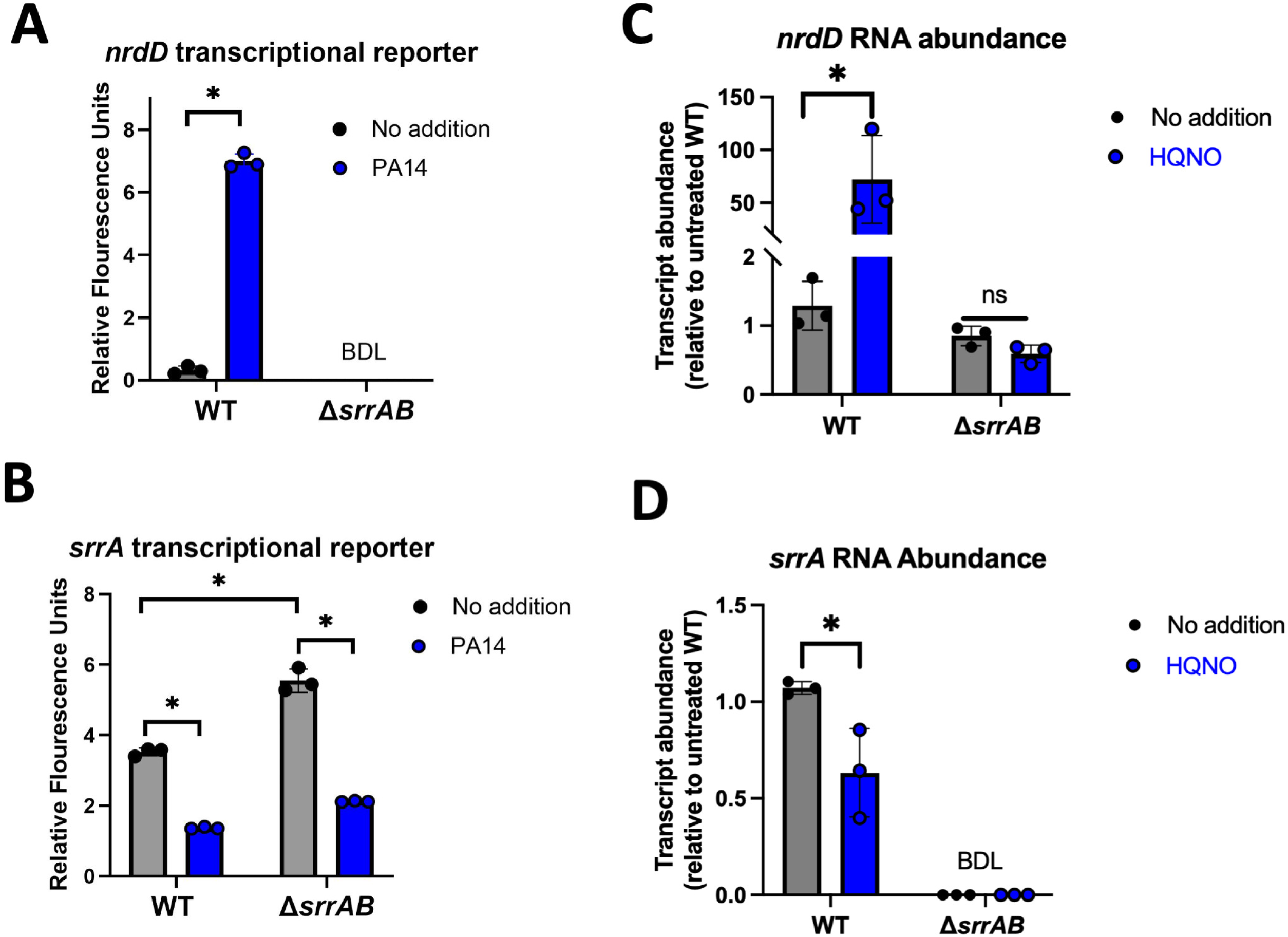
Effect of wild type *P. aeruginosa* cell-free culture spent medium on SrrAB activity and *srrA* transcription. **Panels A and B.** Overnight cultures were diluted into fresh TSB-Cm and challenged with spent media for 8 hours before *gfp* expression was quantified. The expression of *gfp* was quantified in the wild type (WT; JMB1100) and Δ*srrAB* (JMB1467) strains containing the pOS_*nrdD_gfp* (panel A) or pOS_*srrA_gfp* (Panel B) transcriptional reporter after culture in TSB-Cm with or without cell-free spent culture medium isolated from *P. aeruginosa* (PA14). **Panels C and D.** The abundances of mRNA transcripts corresponding to *nrdD* and *srrA* were quantified in the wild type (WT; JMB1100) and Δ*srrAB* (JMB1467) strains using real-time quantitative PCR (qPCR) after culture in TSB with or without cell-free spent culture medium isolated from *P. aeruginosa* (PA14). Data represented the average of biological triplicates with standard deviations shown. Student’s t-tests were performed on the data and * indicates p < 0.05. BDL denotes below the detectable limit.

The *srrA* and *srrB* genes are coded in an operon (*srrAB*) and are directly repressed by SrrA (21). We examined the effect of PA14 spent medium on the transcriptional activity of *srrA*. Spent medium from PA14 significantly decreased *srrA* transcriptional activity in the WT (Fig 2B). The transcriptional activity of *srrA* was increased in the *srrAB* mutant, but *srrA* transcription was also decreased by the spent medium in this strain, suggesting that a transcription factor other than SrrAB was responsible for repressing *srrA* transcription. These findings demonstrate that one or more *P. aeruginosa*-secreted secondary metabolites impact the output of the SrrAB TCRS and at least one additional transcriptional regulator.

To validate the data generated using transcriptional reporters, we performed quantitative real-time PCR (qPCR) to measure the abundance of the *nrdD* and *srrA* transcripts. The *nrdD* transcript was significantly increased when challenged with PA14 spent medium, while the *srrA* transcript showed a significant decrease (Fig. 2C & 2D).

We sought to determine which *P. aeruginosa* secondary metabolites were altering SrrAB regulatory output and *srrA* transcriptional activity. We harvested spent culture media from isogenic *P. aeruginosa* mutants that are deficient in producing one or more secondary metabolites. We then individually added these spent media to *S. aureus* and examined the transcriptional activities of *nrdD* or *srrA*. The spent media from PA14, or mutants defective in either phenazine (Δ*phz*) or hydrogen cyanide (Δ*hcn*) production, increased *nrdD* transcriptional activity, suggesting that the production of these metabolites is not, for the most part, responsible for the altered SrrAB output (Fig 3A). The spent culture medium harvested from the Δ*pqsABC* mutant failed to alter *nrdD* transcriptional activity (Fig 3A). Spent culture media from the PA14, Δ*phz,* or Δ*hcn* strains significantly decreased *srrA* promoter activity, whereas the spent medium from the Δ*pqsABC* mutant did not (Fig 3B). The Δ*srrAB* mutant also responded to the spent media from the PA14, Δ*phz,* or Δ*hcn* strains, reinforcing the hypothesis that the PQS system produces one or more molecules that alter *srrAB* transcription independent of SrrAB.

**Figure 3.**
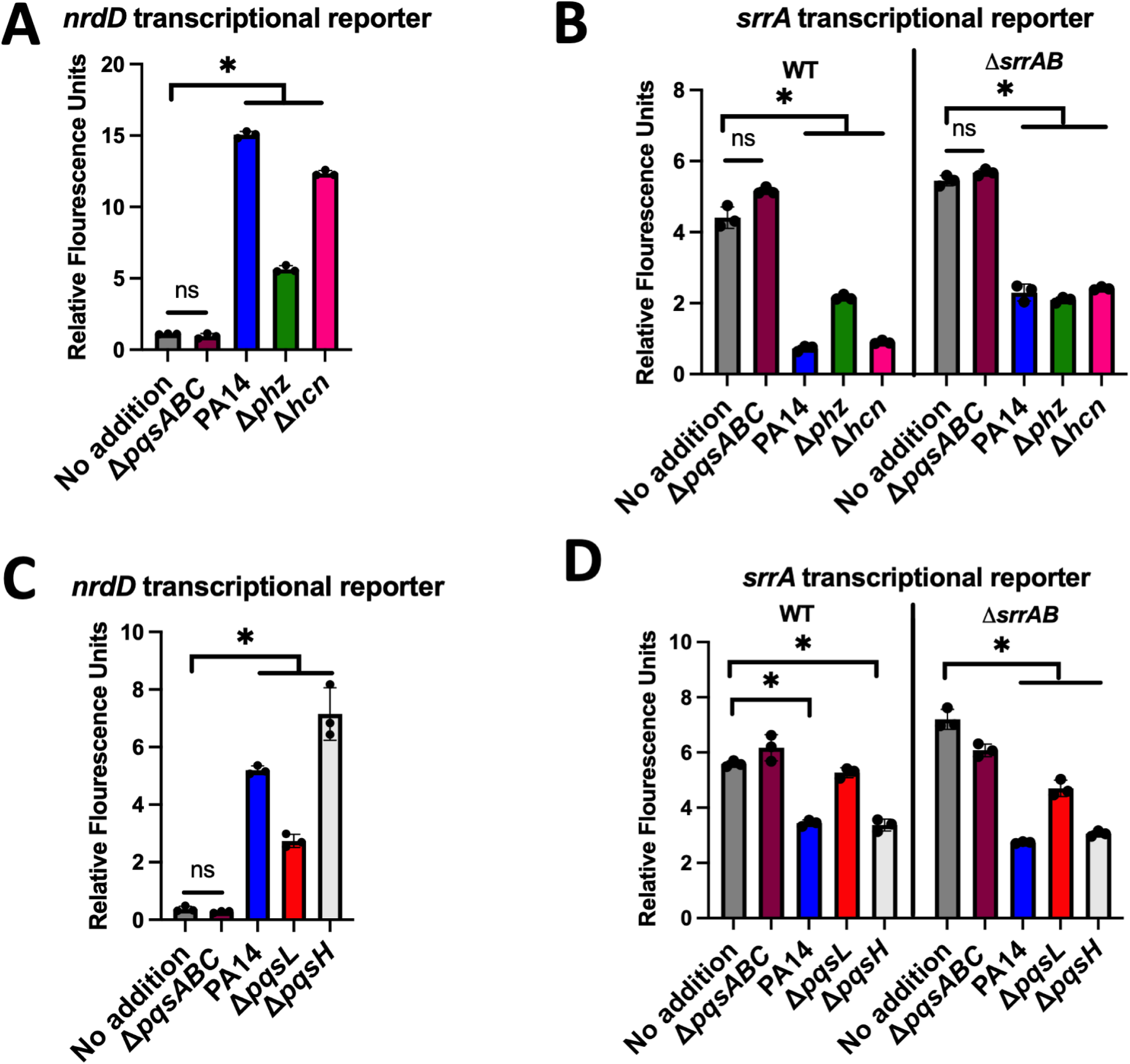
Effects cell-free culture spent media isolated from different *P. aeruginosa* strains on SrrAB activity and *srrA* transcription. The expression of *gfp* was quantified in the wild type and Δ*srrAB* strains containing the pOS_*nrdD_gfp* (panels A and C) or pOS_*srrA_gfp* (Panel B and D) transcriptional reporters after culture in TSB-Cm with or without cell-free spent culture media isolated from various isogenic *P. aeruginosa* strains. Each individual data bar corresponds to the average of the biological replicates, and standard deviations are shown, but the data points sometimes obscure them. Student’s t-tests were performed, and * denoting a p-value ≤ 0.05, and ns denotes no significant difference. BDL denotes below the detectable limit.

We sought to determine which secondary metabolites generated from anthranilic acid were causing changes in *srrA* and *nrdD* transcription. The *pqsL* or *pqsH* gene products are responsible for the production of HQNO and PQS, respectively (Fig 1). We generated spent culture media from isogenic strains incapable of generating one or more anthranilic acid-derived metabolites. Spent media from PA14, Δ*pqsL*, or Δ*pqsH* strains significantly increased *nrdD* transcriptional activity, whereas the spent medium from the Δ*pqsABC* mutant did not (Fig 3C). The spent medium from the Δ*pqsL* had a decreased ability to simulate *nrdD* transcription compared to PA14 spent medium. The spent medium from the Δ*pqsH* mutant phenocopied that seen with PA14 medium.

The spent culture medium from the Δ*pqsL* mutant did not significantly decrease *srrA* transcriptional activity in the WT but did in the Δ*srrAB* mutant (Fig 3D). Spent medium from the Δ*pqsH* strain decreased *srrA* transcriptional activity to a level like that seen with the PA14 spent medium. These results led to a model wherein *P. aeruginosa* produced HQNO alters SrrAB output and alters the activity of an additional regulatory protein that represses *srrA* transcription.

### HQNO stimulates SrrAB and leads to altered *srrA* transcriptional activity

We tested the hypothesis that HQNO stimulates SrrAB and alters the transcriptional activity of *srrA*. We titrated pure HQNO into liquid cultures of the WT containing the *nrdD* transcriptional reporter and observed a significant dose-dependent increase in promoter activity (Fig 4A). The increase in transcriptional activity plateaued at approximately 5 μg mL^-1^ HQNO. We next titrated HQNO into cultures of WT containing the *srrA* transcriptional reporter, which resulted in a dose-dependent decrease in *srrA* transcriptional activity, and again, the maximal effect occurred at approximately 5 μg mL^-1^ (Fig 4B).

**Figure 4.**
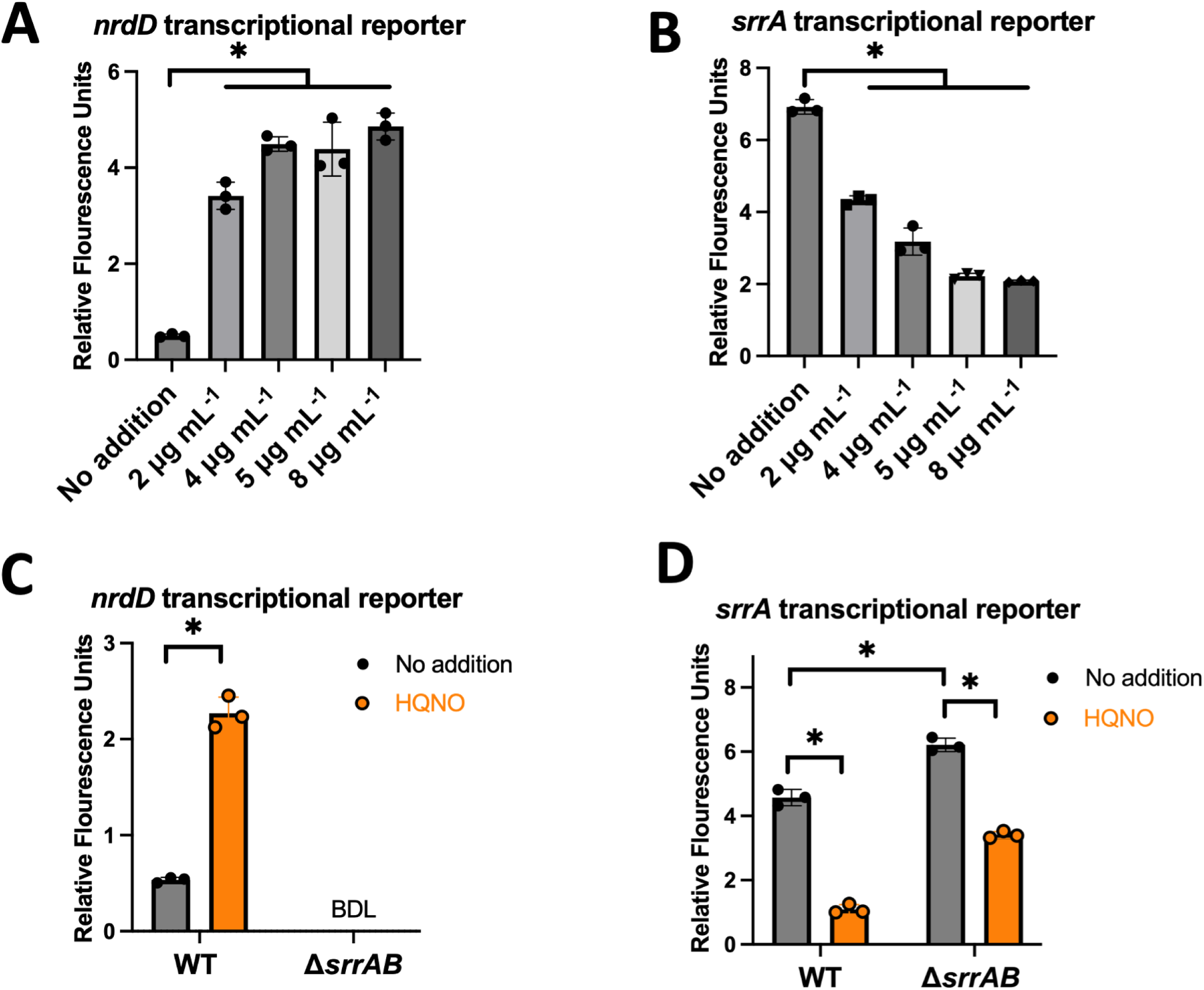
HQNO alters SrrAB activity and *srrA* transcription. **Panels A and B.** The wild type (WT; JMB1100) strain containing the pOS_*nrd_gfp* (panel A) or pOS_*srrA_gfp* (panel B) transcriptional reporters were co-cultured with different concentrations of HQNO before *gfp* expression was quantified using fluorimetry. **Panels C and D.** The WT and Δ*srrAB* (JMB1467) strains containing the pOS_*nrdD_gfp* (panel C) or pOS_*srrA_gfp* (Panel D) transcriptional reporter trains were cultured in TSB-Cm with and without 5 μg mL^-1^ HQNO before *gfp* expression was quantified using fluorimetry. Each individual data bar corresponds to the average of the biological replicates, and standard deviations are shown, but the data points sometimes obscure them. Student’s t-tests were performed and * denoting a p-value ≤ 0.05. BDL denotes below the detectable limit of the fluorimeter.

We sought to determine if the observed transcriptional effects of HQNO on *srrA* and *nrdD* require SrrAB (Fig 4C-D). The transcriptional activity of *nrdD* did not increase in the Δ*srrAB* strain upon treatment with HQNO, demonstrating a dependency on SrrAB (Fig 4C). Consistent with previous results, the effect of HQNO on *srrA* transcriptional activity was independent of SrrAB (Fig 4D). The data presented thus far are consistent with a model wherein *P. aeruginosa* secreted HQNO alters SrrAB activity and *srrA* transcriptional activity, which occurs through an alternate transcriptional regulator.

### Proficient respiration is required for HQNO to alter *srrA* transcription

It has been hypothesized that HQNO inhibits the cytochrome oxidases of *S. aureus,* resulting in the inhibition of respiration (9, 26). We tested the hypothesis that HQNO inhibited *S. aureus* respiration under the growth conditions used for our study. *S. aureus* utilizes two terminal oxidases called Cyd and Qox (27). We monitored oxygen consumption in the WT and a strain lacking both terminal oxidases, after culture in the presence or absence of HQNO. The addition of HQNO significantly decreased the rate of dioxygen consumption in the WT (Fig 5A). In contrast, no dioxygen consumption was noted for the Δ*cydA qoxB::tet* strain, and the presence of HQNO did not alter this phenotype.

**Figure 5.**
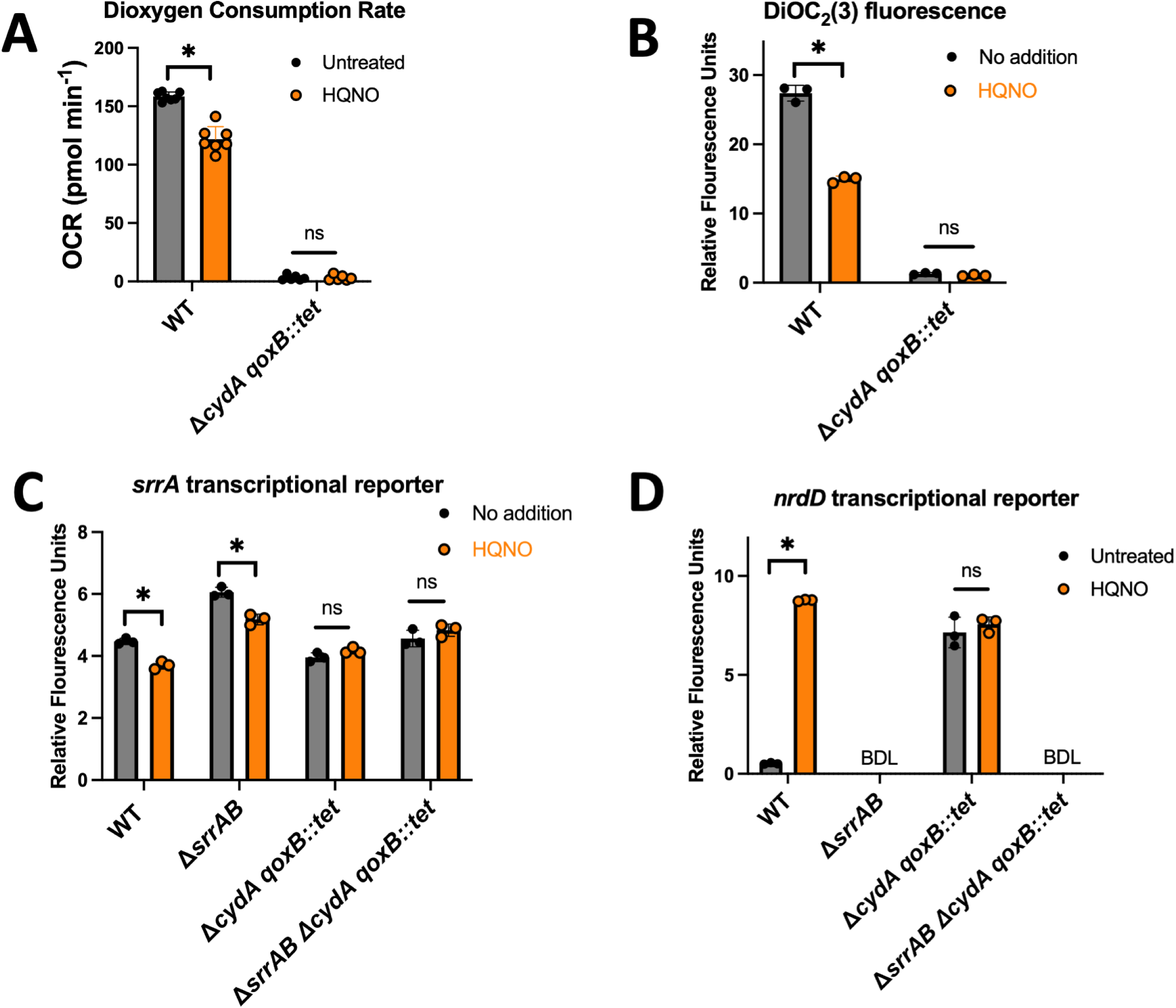
HQNO and dioxygen respiration. **Panel A.** Oxygen consumption rates were quantified in the wild type (WT; JMB1100) and Δ*cydA qoxB::tet* (JMB 8988) strains after culture in TSB media with the presence or absence of 5 μg mL^-1^ HQNO. **Panel B.** Membrane potential was qualitatively assessed using 1 mM of the fluorescent dye DIOC_2_(3) in the WT and Δ*cydA qoxB*:*:tet* strains after culture in TSB media with the presence or absence of 5 μg mL^-1^ HQNO. **Panels C and D.** The WT, Δ*srrAB* (JMB1467), Δ*cydA qoxB*:*:tet*, and Δ*srrAB* Δ*cydA qoxB*:*:tet* (JMB 8989) strains containing pOS_*nrdD_gfp* or pOS_*srrA_gfp* after culture in TSB media in the presence and absence of 5 μg mL^-1^ HQNO before *gfp* expression was quantified using fluorimetry. Each individual data bar corresponds to the average of the biological replicates, and standard deviations are shown, but the data points sometimes obscure them. Student’s t-tests were performed, and * denoting a p-value ≤ 0.05, and ns denotes no significant difference. BDL denotes below the detectable limit.

We tested the hypothesis that challenging *S. aureus* with HQNO would decrease the membrane potential, which is a respiration byproduct. The fluorescent dye DiOC_2_(3) can be used to qualitatively compare the membrane potential of bacterial strains (28, 29). DiOC_2_(3) is positively charged and accumulates in a charge-dependent manner in the cells and shifts from green to red as the membrane potential increases (30). We treated aerobically cultured WT and Δ*cydA qoxB::tet* strains with HQNO and quantified membrane potential. DiOC_2_(3) attributed red fluorescence was lower in the Δ*cydA qoxB::tet* mutant when compared to that of WT (Fig 5B). The addition of HQNO decreased red fluorescence in the WT but did not alter the fluorescence of the Δ*cydA qoxB::tet* strain.

We next tested the hypothesis that the alterations in the transcriptional activities of *nrd* and *srrA* upon HQNO addition required the cells to be respiring. We compared *srrA* transcriptional activity in the WT and Δ*cydA qoxB::tet* strains and their isogenic Δ*srrAB* mutants after aerobic culture with HQNO. We again noted that transcriptional activity decreased in the WT and Δ*srrAB* strains cultured with HQNO (Fig 5C). Transcriptional activity of *srrA* was decreased in the Δ*cydA qoxB::tet* strain compared to the WT, but there was no change in transcriptional activity up growth with HQNO. The transcriptional activity of *nrdD* was increased in the Δ*cydA qoxB::tet* strain, consistent with *nrdD* transcription being controlled by respiratory status, and transcriptional activity was not affected by HQNO in the Δ*cydA qoxB::tet* strain (Fig 5D). These data varify that HQNO inhibits *S. aureus* dioxygen respiration and demonstrate that cells must be respiring for HQNO to alter SrrAB output and alter *srrA* transcription.

### Growth with HQNO alters metabolite pools, resulting in Rex repression of *srrAB* transcription

We tested the hypothesis that co-culture with HQNO inhibits cellular respiration, which alters metabolite pools, modulating the activity of a transcriptional regulator that modifies *srrA* transcription. We conducted an untargeted metabolomic analysis to identify metabolites that have altered abundances upon co-culture with HQNO.

We identified several metabolites that were either significantly increased or decreased upon co-culture with HQNO (Tables 1 and S1). There were significant increases in NAD^+^, lactate, pyruvate, and acetyl-CoA, which correlates with *S. aureus* increasing fermentation and decreasing TCA cycle activity to balance redox upon HQNO inhibition of dioxygen respiration. The concentrations of the amino acids alanine, glycine, threonine, and proline were significantly increased. Alanine can be synthesized from pyruvate via alanine dehydrogenases (27). Glycine is synthesized from threonine and or serine, which also showed an increase with HQNO.

The metabolites that had the largest decrease upon co-culture with HQNO are intermediates in arginine synthesis or utilized in nitrogen metabolism, including citrulline, aspartate, and glutamate (Tables 1 and S1). The one exception was ornithine, which was significantly increased with HQNO and could explain the increase in proline as this amino acid is synthesized from ornithine (31). An additional explanation for these findings is the altered regulation of the *arcADRBC* operon, which codes for the arginine deiminase system. Arc functions in energy generation during non-respiratory growth by importing arginine using an arginine ornithine antiporter and then converting arginine to ornithine in the cytosol, generating ATP (32, 33).

NAD^+^, but not NADH, increased upon HQNO treatment, resulting in an increase in the NAD^+^/NADH ratio (Fig 6A and Table 2). Likewise, there was an overall increase in oxidized nicotinamide adenine dinucleotide pools compared to their reduced counterparts (Table 2). Accumulation of NAD^+^ increases the affinity of the *S. aureus* transcriptional regulator Rex for DNA *in vitro* (34, 35). Rex controls the transcription of the *arc* operon, and there is a Rex binding site in the promoter of *srrA* that ends -85 base pairs from the translational start site (34). Rex directly associates with the *srrA* operator *in vitro* (34, 35).

**Figure 6.**
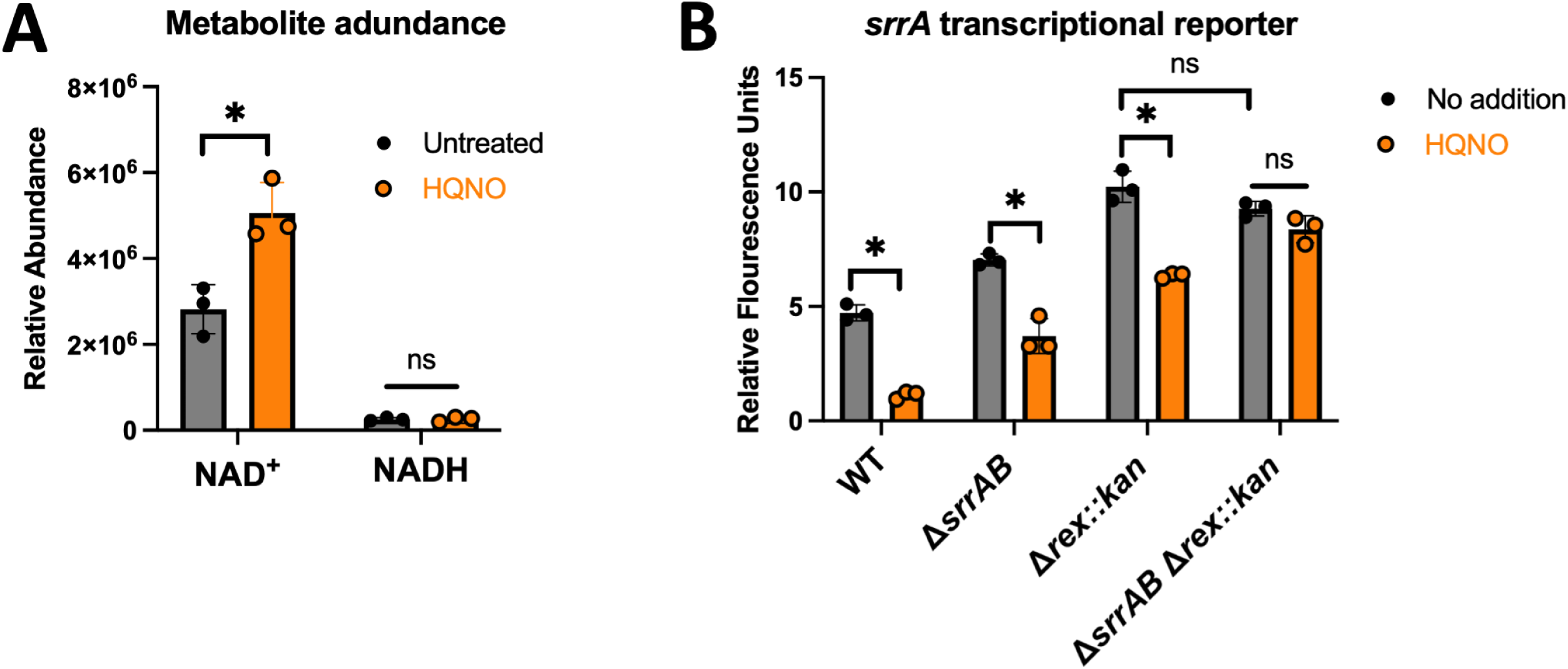
Rex represses *srrA* transcription in response to HQNO challenge. **Panel A.** The relative abundances of NAD^+^ and NADH in the wild type (WT; JMB1100) strain after co-culture in the presence and absence of 5 μg mL^-1^ HQNO. **Panel B.** The expression of *gfp* was monitored in the WT, Δ*srrAB* (JMB1467), Δ*rex::kan* (JMB 13604), and Δ*srrAB* Δ*rex::kan* (JMB 13626) strains containing pOS_*srrA_gfp* after culture in the presence and absence of 5 μg mL^-1^ HQNO. The response to HQNO requires both SrrAB and Rex regulators. Each individual data bar corresponds to the average of the biological triplicates, and standard deviations are shown, but the data points sometimes obscure them. Student’s t-tests were performed, and * denoting a p-value ≤ 0.05, and ns denotes no significant difference. BDL denotes below the detectable limit.

We tested the hypothesis that Rex was responding to growth in the presence of HQNO to repress *srrA* transcription. We quantified *srrA* transcriptional activity in the WT, Δ*rex*, Δ*srrAB*, and Δ*rex* Δ*srrAB* strains after growth in the presence or absence of HQNO. Transcriptional activity of *srrA* was increased in the Δ*rex,* Δ*srrAB*, and Δ*rex* Δ*srrAB* strains when compared to the WT (Fig. 6B). The phenotype of the Δ*rex* mutation was dominant and the Δ*rex* Δ*srrAB* phenocopied the Δ*rex* strain. The addition of HQNO decreased *srrA* transcriptional activity in the WT, Δ*rex,* and Δ*srrAB* strains but not in the Δ*rex* Δ*srrAB* strain. These findings demonstrate that Rex and SrrA respond to growth in the presence of HQNO and independently repress *srrA* transcription.

### SrrAB contributes to fitness and survival when *S. aureus* is exposed to HQNO or PA14

We monitored growth kinetics to test the hypothesis that SrrAB and Rex promote fitness when challenged with HQNO. The WT, Δ*srrAB*, Δ*rex*, and Δ*srrAB* Δ*rex* mutants all had lower growth rates in TSB containing HQNO, but the growth rate of the Δ*srrAB* mutant was lower than that of the WT, whereas the Δ*rex* mutant was not (Fig 7A). The growth rate of the Δ*rex* Δ*srrAB* double mutant phenocopied that of the Δ*srrAB* strain. Expressing *srrAB* from the native promoter genetically complemented the HQNO-challenged growth rate phenotype of the Δ*srrAB* mutant (Fig 7B). These findings demonstrate that SrrAB is needed to maintain fitness when exposed to HQNO.

**Figure 7.**
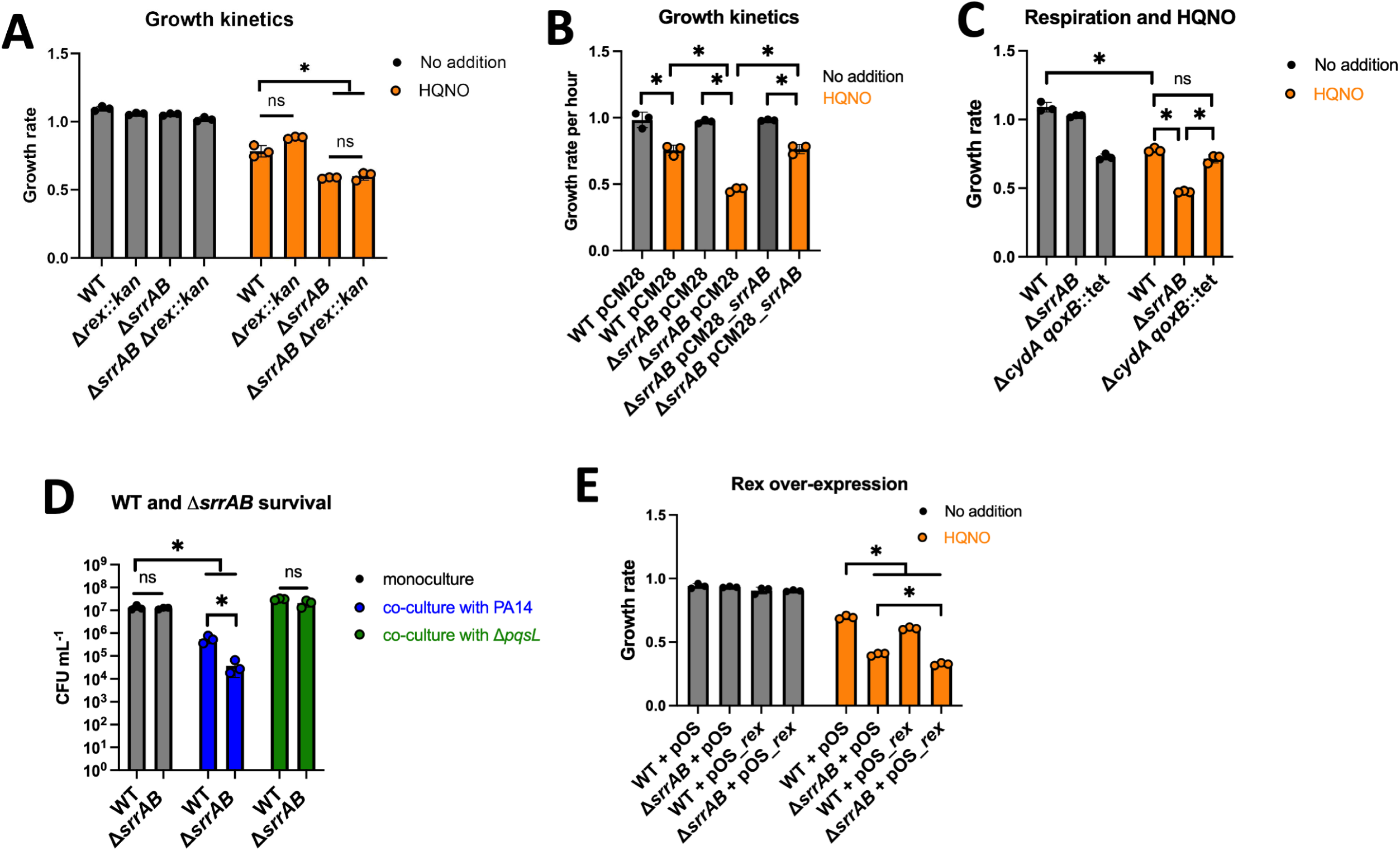
SrrAB is required for fitness and survival when challenged with HQNO and *P. aeruginosa*. **Panel A.** The growth rates for the WT, Δ*srrAB* (JMB1467), Δ*rex::kan* (JMB 13604), and Δ*srrAB* Δ*rex::kan* (JMB 13626) strains were quantified after growth in TSB media with or without 5 μg mL^-1^ of HQNO. **Panel B.** The growth rates of the WT or Δ*srrAB* strains containing pCM28 or pCM28_*srrAB* were quantified after growth in in TSB media with or without 5 μg mL^-1^ of HQNO. **Panel C.** The growth rates for the wild type (WT; JMB1100), Δ*srrAB* (JMB1467), and Δ*cydA qoxB::tet* (JMB8988) strains were quantified after growth in TSB with and without 5 μg mL^-1^ of HQNO. **Panel D.** *P. aeruginosa* PA14 (JMB 10389) or Δ*pqsL* (JMB13827) strains were co-cultured with the *S. aureus* WT or Δ*srrAB* mutant before *S. aureus* survival was determined by counting colony-forming units (CFU) after plating on TSB with 7.5% NaCl. **Panel E**. The growth rates of the WT and Δ*srrAB* containing either pOS1 or pOS1_*rex* were quantified after culture in TSB-Cm media with and without 5 μg mL^-1^ of HQNO. Each individual data bar corresponds to the average of the biological triplicate, and standard deviations are shown, but the data points sometimes obscure them. Student’s t-tests were performed, and * denoting a p-value ≤ 0.05, and ns denotes no significant difference.

SrrAB is a positive regulator of respiration and fermentation when respiration is impaired (36). We further investigated the role of SrrAB in improving fitness and survival when challenged with HQNO. We aerobically cultured the WT, Δ*srrAB*, and Δ*cydA qoxB*::*tet* strains in the presence and absence of HQNO. The Δ*cydA qoxB*::*tet* strain had a lower growth rate than the WT strain, which HQNO did not alter, consistent with HQNO inhibiting respiration (Fig 7C). The growth rates of the WT and Δ*cydA qoxB*::*tet* strains were similar when challenged with HQNO, whereas the growth rate was lower for the Δ*srrAB* mutant. These data are consistent with a model wherein the role of SrrAB in increasing fitness when challenged with HQNO is more than just increasing the transcription of genes coding respiratory components.

*P. aeruginosa* can outcompete *S. aureus* when they are co-cultured. The finding that SrrAB was required for fitness of HQNO-challenged *S. aureus* led to the hypothesis that SrrAB is required for survival when co-cultured with *P. aeruginosa*. We quantified the survival of the WT and Δ*srrAB* strains with and without co-culture with strains of *P. aeruginosa* that can and cannot produce HQNO. The Δ*srrAB* mutant had a lower survival than the WT when co-cultured with PA14 (Fig 7D). The killing of the WT and Δ*srrAB* mutant was abrogated upon co-culture with the *P. aeruginosa* Δ*pqsL* mutant, demonstrating that HQNO was a primary determinant that allowed *P. aeruginosa* to outcompete *S. aureus* in co-culture under the conditions tested. These data demonstrate that SrrAB increases the ability of *S. aureus* to compete with an HQNO-producing strain of PA14.

Overexpression of Rex negatively impacts *S. aureus* growth upon respiration inhibition because it represses the transcription of genes utilized in fermentation, including those coding lactate dehydrogenases (34). We created *S. aureus* strains with a plasmid containing *rex* under the transcriptional control of the constitutive *lgt* promoter (pOS_*rex*). The WT and Δ*srrAB* strains containing pOS_*rex* had a lower growth rate than those containing the empty vector (pOS) upon HQNO challenge (Fig 7C). These data indicate that increasing the expression of *rex* decreases the ability of *S. aureus* to overcome HQNO challenge.

### Effective fermentation promotes fitness and survival when *S. aureus* is challenged with HQNO or PA14

We next tested the hypothesis that effective fermentation is required for fitness upon HQNO challenge. SrrAB regulates the genes coding lactate dehydrogenases and pyruvate formate lyase (Pfl) (24, 25, 34). During fermentative growth, Pfl is utilized to produce acetyl-CoA (37). We determined that the growth rate of a *pflB::Tn* mutant was significantly decreased compared to the WT upon co-culture with HQNO (Fig. 8A).

**Figure 8.**
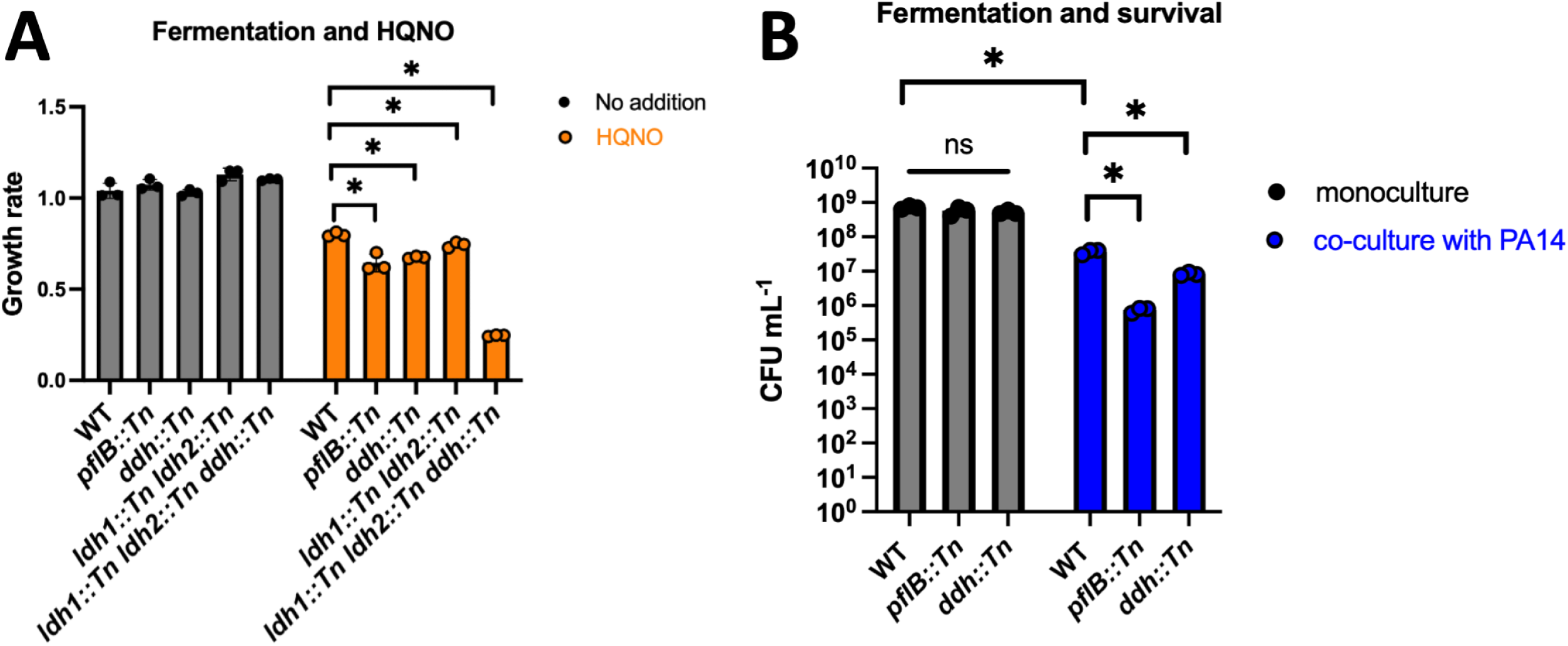
Effective fermentation is important for *S. aureus* fitness and survival upon HQNO or *P. aeruginosa*. **Panel A.** The growth rates of the WT, *pfl::Tn* (JMB 10737), *ddh::Tn* (JMB 14930), *ldh1::Tn ldh2::Tn* (JMB 14175), and *ldh1::Tn ldh2::Tn ddh::Tn* (JMB 14176) strains were quantified after growth in TSB media with and without 5 μg mL^-1^ of HQNO. **Panel B.** *P. aeruginosa* PA14 (JMB 10389) was co-cultured with the *S. aureus* WT, *pflB::Tn*, or *ddh::Tn* strains before *S. aureus* survival was determined by counting colony-forming units (CFU) after plating on TSB with 7.5% NaCl. Each individual data bar corresponds to the average of the biological replicates, and standard deviations are shown, but the data points sometimes obscure them. Student’s t-tests were performed, and * denoting a p-value ≤ 0.05, and ns denotes no significant difference.

Lactate, generated from pyruvate, is a primary *S. aureus* fermentation byproduct and lactate accumulated upon treatment with HQNO (Table 1). The *S. aureus* genome codes for two cytosolic L-lactate dehydrogenases (*ldh1*, *ldh2*), a cytosolic D-lactate dehydrogenase (*ddh*), and a quinone-linked lactate dehydrogenase (*lqo*) (38). We examined the growth kinetics of the WT, *ldh1::Tn ldh2::Tn*, and *ddh::Tn* strains with and without HQNO. All strains had a significant decrease in growth rate when exposed to HQNO. The growth rates of the *ldh1::Tn ldh2::Tn* double mutant and *ddh::Tn* strains were lower than that of the WT, with the *ddh::Tn* mutant having a stronger phenotype (Fig. 8A). The phenotypes associated with the *ldh1::Tn ldh2::Tn* and *ddh::Tn* mutations had an additive effect upon HQNO-challenge. These data highlight the importance of using lactate too balance redox upon HQNO-dependent inhibition of respiration.

**Table 1.** Metabolites that are significantly altered in *S. aureus* upon growth with HQNO.

| **Metabolite** | **Log2 fold-change from non-treated** |
| --- | --- |
| Citrulline | -5.6 |
| NADPH | -3.9 |
| Arginine | -2.2 |
| Aspartate | -2.2 |
| N-Acetylaspartate | -1.8 |
| NADP<sup>+</sup> | -1.4 |
| Glutamate | -1.4 |
| Coenzyme A | -1.3 |
| Pyruvate | 0.7 |
| NAD<sup>+</sup> | 0.8 |
| Acetyl-CoA | 1.0 |
| Alanine | 1.6 |
| Ornithine | 2.1 |
| Glycine | 2.4 |
| Proline | 2.6 |
| Lactate | 2.9 |
| Threonine | 3.9 |

**Table 2.** Ratios for nicotinamides when HQNO is present.

| **Table 2. Ratios for nicotinamides when HQNO is present.** |  |  |
| --- | --- | --- |
| **Metabolites** | No addition | HQNO |
| NAD<sup>+</sup> / NADH | 10.9 | 19.4 |
| NAD<sup>+</sup> + NADP<sup>+</sup> / NADH + NADPH | 6.2 | 19.2 |

For a full list of metabolites, refer to Supplemental Table 1.

We hypothesized that the survival of *S. aureus* strains defective in fermentation would be decreased upon co-culturing with *P. aeruginosa*. We quantified the survival of the WT, *pflB::Tn*, and *ddh::Tn* strains with and without co-culture with PA14. Both the fermentation-deficient strains had decreased survival when compared to the WT (Fig 8B). The findings demonstrate that effective fermentation is required for fitness and survival when challenged with *P. aeruginosa*.

## Discussion

*P. aeruginosa* secretes secondary metabolites that negatively impact the fitness of *S. aureus* (13, 15, 39, 40). The decrease in fitness ultimately leads *P. aeruginosa* domination in co-cultures and *S. aureus* death. This initial study aimed to determine if *S. aureus* senses the presence of *P. aeruginosa* secondary metabolites and, if so, determine whether the regulatory response improves *S. aureus* fitness in co-culture. In *S. aureus*, SrrAB increases the transcription of genes utilized for fermentation and respiration when the respiration rate decreases (22, 23, 25). The results herein demonstrate that the SrrAB TCRS senses and responds to the presence of *P. aeruginosa* spent medium containing secondary metabolites. We used *P. aeruginosa* mutant strains and pure compound to determine that the PQS quorum sensing system-derived molecule HQNO was responsible for stimulating SrrAB. Previous studies have demonstrated that during co-culture *P. aeruginosa* produced HQNO drives *S. aureus* towards a fermentative phenotype (17). Importantly, HQNO has been detected in cystic fibrosis patients infected with *P. aeruginosa,* and our results demonstrate that SrrAB promotes *S. aureus* survival when these two bacteria are co-cultured in the laboratory (16, 41).

Previous work in our lab determined that either chemical or genetic inhibition of respiration increased SrrAB activity (21). Herein, we determined that treatment of WT *S. aureus* with HQNO, but not a mutant lacking both terminal oxidases, resulted in decreased dioxygen consumption, a decreased proton motive force, and a decreased growth rate. The transcriptional activities of *srrA* and *nrdD* are regulated by SrrAB, and the transcriptional activities of these promoters upon treatment with HQNO mimicked the transcriptional phenotypes of the mutant strain lacking the terminal oxidases. These data support the working model in Figure 9, where *P. aeruginosa*-generated HQNO inhibits *S. aureus* respiration, which is sensed by SrrAB and responds by altering the transcription of genes utilized for fermentation. Consistent with this model, previous studies discovered that the co-culture of *P. aeruginosa* and *S. aureus* increased the transcription of SrrAB-regulated genes (28, 35).

**Figure 9.**
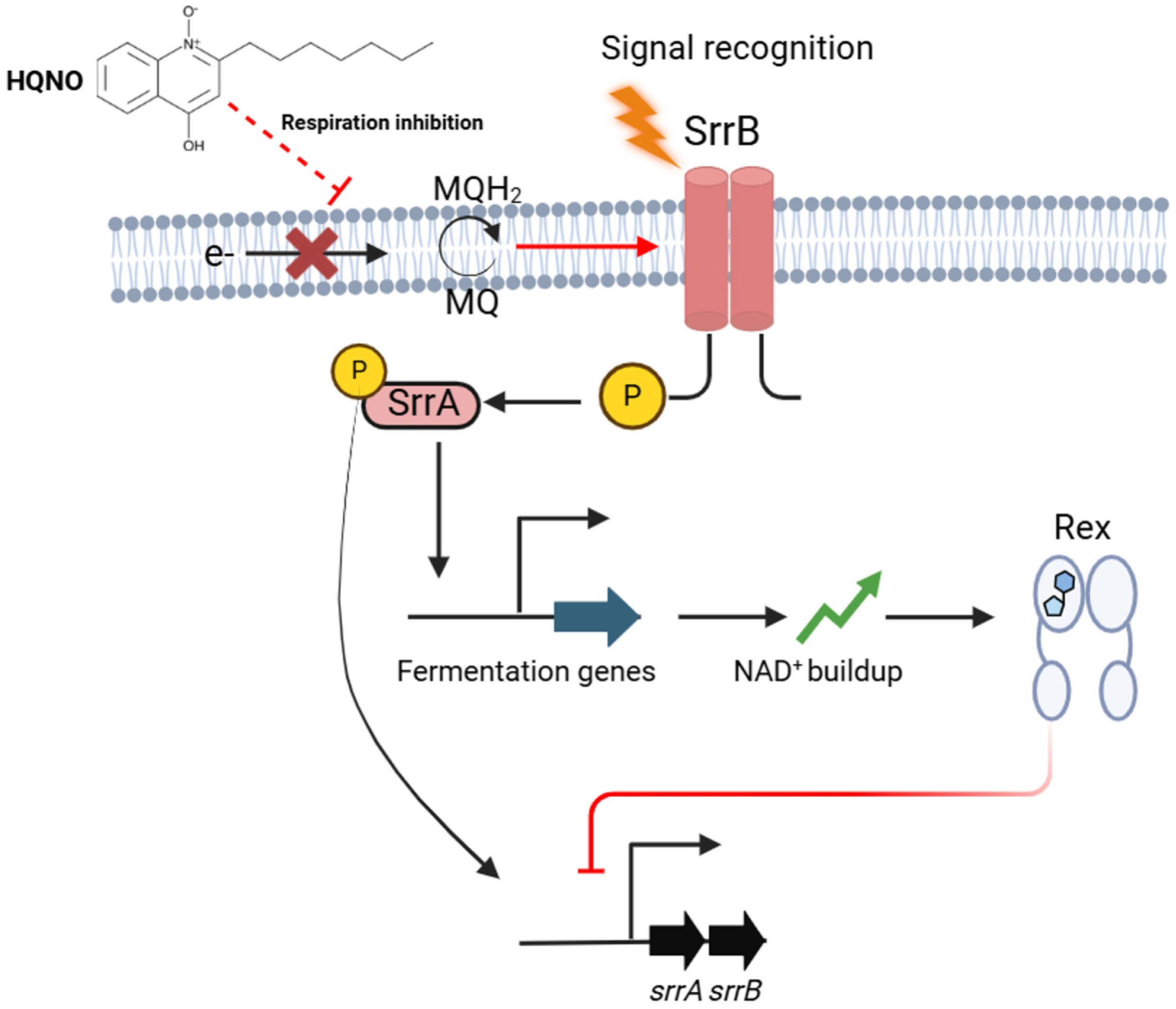
Proposed model for SrrAB regulation by Rex. *P. aeruginosa*-produced. HQNO inhibits dioxygen respiration in *S. aureus,* stimulating the histidine kinase SrrB, which alters the titers of phosphorylated SrrA. SrrA then represses *srrAB* transcription and increases the transcription of genes utilized for fermentative and anaerobic growth. This exacerbates a fermentation phenotype and increases the ratio of NAD^+^ to NADH. The increased abundance of NAD^+^ binds to Rex, increasing its affinity to the promoter of *srrAB* and repressing its transcription.

HQNO is structurally similar to menaquinone, which is the lipid-soluble electron carrier that *S. aureus* utilizes to shuttle electrons between respiratory components (42). It has been hypothesized that one mechanism for HQNO-dependent inhibition of *S. aureus* respiration is by competing with menaquinone for binding to NADH:oxidoreductase (NDH-2). HQNO decreased the rate of *S. aureus* NDH-2 reoxidation after NADH-dependent FAD reduction with a K_i_ app value of 6.8 ± 0.4 μM (43). The *Caldalkalibacillus thermarum* NDH-2 was inhibited with IC_50_ of 7.3 μM in the presence of 50 μM menadione, which is a water-soluble menaquinone analog (44). Challenge of *S. aureus* with HQNO has been linked to increased biofilm formation, a trait of anaerobic cultured *S. aureus* (45, 46). We chose to work with a strain lacking the terminal oxidases because there are numerous electron inputs to the respiratory chain, including succinate dehydrogenase, malate quinone oxidoreductase, and a quinone-linked lactate dehydrogenase, but only two enzymes that can transfer these electrons to dioxygen. Further biochemical analyses will be required to determine the precise respiratory component(s) inhibited by HQNO.

To our surprise, the transcriptional activity of *srrA* was altered in a Δ*srrAB* mutant in response to HQNO treatment. Three findings led to the hypothesis that Rex was responsible for this phenotype. First, metabolomic analyses found that the concentrations of metabolites associated with nitrogen metabolism were the most altered upon HQNO treatment. The concentrations of citrulline and arginine were decreased while ornithine increased. These data suggested altered regulation of the *arcABCD* operon, which functions in energy generation during non-respiratory growth. The *arcABCD* operon is directly regulated by Rex (35). Second, HQNO treatment increased the NAD^+^ / NADH ratio and increasing this ratio increases the DNA binding activity of Rex (35). Third, work by others determined that Rex associates with the *srrA* operator, and *srrA* transcription is increased in a *rex* mutant (34, 35). Consistent with our hypothesis, epistasis experiments presented herein demonstrated that both SrrA and Rex repress *srrA* transcription, and HQNO treatment did not decrease *srrA* transcription in the *srrAB rex* double mutant strain.

When the NAD^+^ / NADH ratio rises, the affinity of Rex for DNA increases, resulting in repressed transcription of genes involved in fermentation, including those coding lactate dehydrogenases, alcohol dehydrogenases, and pyruvate formate lyase (35). Rex also represses *srrAB* transcription, which is likely because SrrAB stimulation increases the transcription of genes involved in fermentation (21, 23). It seems counterintuitive that inhibition of respiration would increase the ratio of NAD+ / NADH, resulting in increased Rex activity; however, the experiments presented examined transcriptional and metabolomic changes that occurred after prolonged exposure to HQNO, providing the necessary time for *S. aureus* to adjust to respiration inhibition. Consistent with this, growth with HQNO increased lactate accumulation, which coincides with past findings demonstrating that coculturing increased the production of lactate and alcohol dehydrogenases (17). Rex and SrrA regulate the expression of *ldh* and *adh* genes (35, 47).

Upon respiratory inhibition, SrrAB stimulation increased transcription genes involved in fermentation (21, 22). Respiratory inhibition also decreases the ratio of NAD^+^ / NADH alleviating Rex repression. We determined that the Δ*srrAB* mutant had decreased growth with HQNO, but the Δ*rex* mutant behaved like the WT. The slow growth phenotype associated with the Δ*srrAB* mutation was exacerbated in a respiration-deficient strain, which is likely because SrrAB is an activator of transcription of genes utilized for fermentation. The Δ*rex* mutant likely behaved like WT because Rex is a repressor, and the transcription of genes that promote fermentation was alleviated upon HQNO treatment in both the WT and Δ*rex* strains. In support of this hypothesis, increased expression of *rex* decreased the growth rate when HQNO was present, which was presumably because of increased Rex repression of genes utilized for fermentative redox balance. This finding correlates with a previous study demonstrating that Rex overexpression decreased *S. aureus* growth when respiration was inhibited by nitric oxide (34).

The finding that the ratio of NAD^+^ /NADH increased suggested increased fermentation upon HQNO treatment. Increased production of lactate dehydrogenases, which oxidize NADH, are a primary mechanism to balance redox during fermentation. The decreased growth rates in the lactate dehydrogenase mutant strains upon HQNO addition demonstrated the importance of redox balance upon HQNO-dependent respiratory inhibition (Fig. 8A).

The results of this study demonstrate that *P. aeruginosa* secreted HQNO inhibits respiration in *S. aureus*. This inhibition is sensed by the SrrAB TCRS, which responds by increasing the transcription of genes involved in fermentation and redox balance. This favors the oxidation of NADH to NAD^+^, activating Rex, which represses *srrAB* transcription. The inability of *S. aureus* to sense respiration inhibition and respond using SrrAB or utilize Pfl or lactate dehydrogenases decreases cell fitness and survival when co-cultured with HQNO or *P. aeruginosa*, respectively. Future studies using infection models and the integration of host physiology are necessary to determine how these findings shed light on *S. aureus* and *P. aeruginosa* competition in patients suffering from cystic fibrosis.

## Material and Methods

### Bacterial strains and experimental conditions

Tryptic soy broth (TSB) purchased from VWR was used for liquid growth analyses. Solid media was supplemented with 1.5% agar (VWR). Unless stated differently, aerobic cultures were grown in 2 mL of TSB in 10 mL culture tubes shaking at 200 rpm at 37 °C. Tubes were slanted at a 30-degree angle to promote aeration. Cells were treated with 5 µg mL^-1^ HQNO (Selleck Chemicals) prepared as a 5 mg mL^-1^ stock in DMSO or vehicle control. Plasmids or chromosomal insertions were selected using antibiotics at a final concentration of 30 µg mL^-1^ chloramphenicol or 10 µg mL^-1^ for erythromycin (Erm), kanamycin (Kan), and tetracycline (Tet). Plasmids were maintained by supplementing the media with 10 µg/mL^-1^ Cm (TSB-Cm). DNA primers were purchased through Integrated DNA Technologies (Coralville, IA).

### Plasmids and construction of bacterial strains

All plasmids and bacterial strains utilized for this study are listed in Table 3. All transductions were performed using bacteriophage 80α (48). All *S. aureus* strains used were isogenic with USA300_LAC (wild type; WT) and erythromycin sensitive (49). Bacterial strains and plasmids were PCR verified and subsequently sequenced by Azenta (South Plainfield, NJ). Amplicons were generated using USA300_LAC DNA as a template and DNA primer pairs (Table 4). Amplicons were digested with *KpnI-*HF and *HindIII-*HF (New England Biolabs) for 1 hour at 37°C followed by heat inactivation at 80°C for 20 min and ligated into similarly digested pOS_*saeP1_gfp* (50) using Quick Ligase (New England Biolabs). The ligation product was transformed into *E. coli* DH5-α and selected for growth in lysogeny broth (LB) agar plates containing 1 mg mL^-1^ ampicillin. Colonies were verified for insert presence by PCR, and plasmids were extracted. Plasmids were transformed into *S. aureus* RN4220 (51), and cells were selected for Cm resistance. Plasmids were then transduced into USA300_LAC background using bacteriophage α80 (48).

**Table 3.**
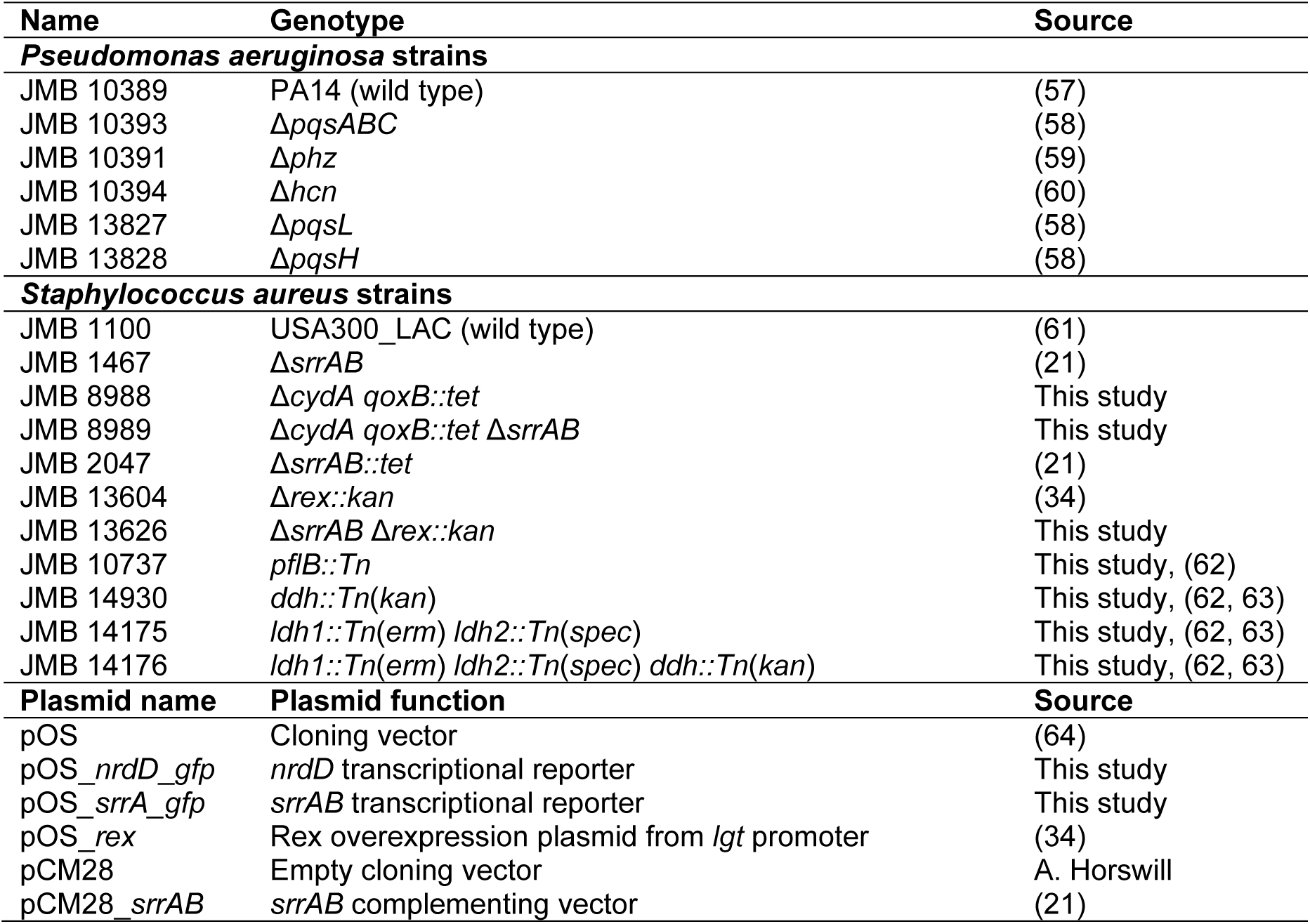
Strains and plasmids used in this study.

**Table 4.**
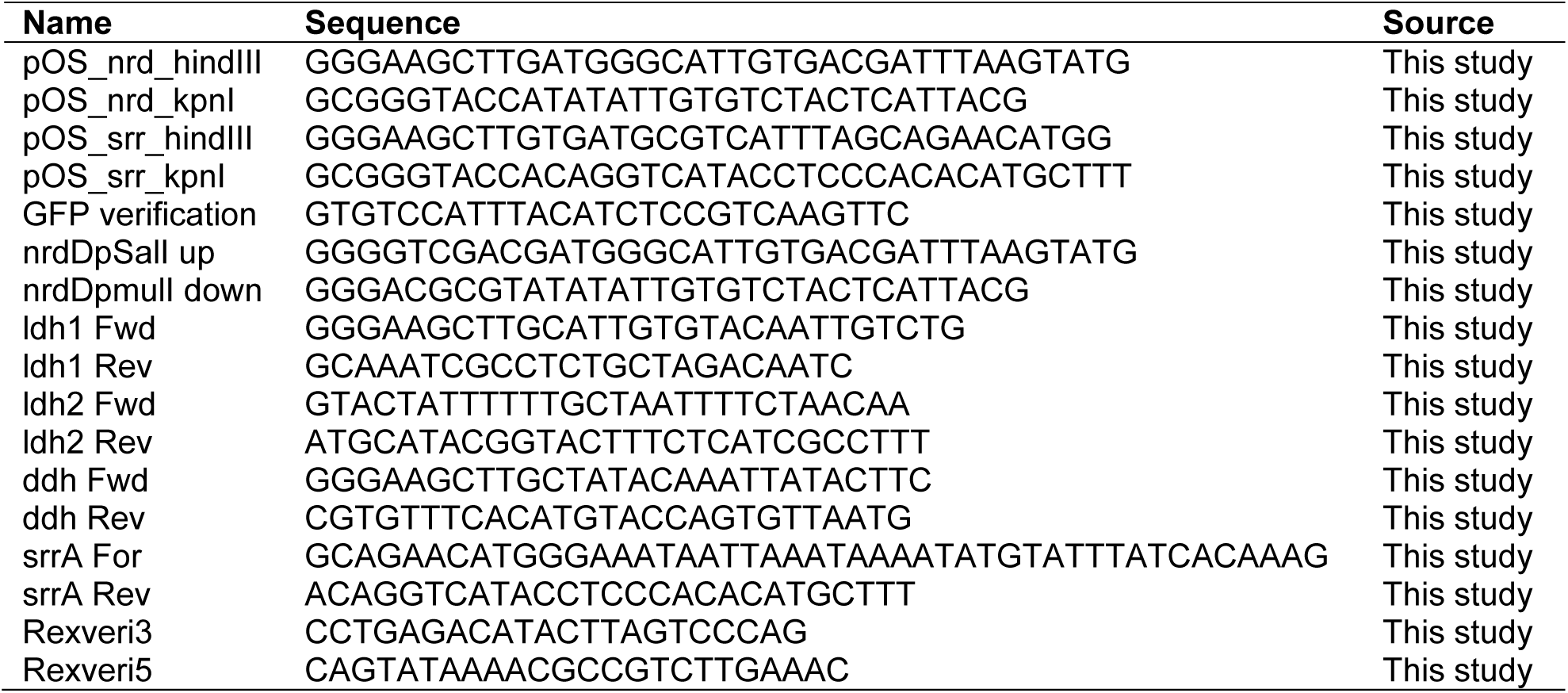
Primers used for this study.

### Kinetic growth analyses

To assess the impact of *P. aeruginosa* on *S. aureus* fitness, we cultured *S. aureus* strains in triplicate overnight in TSB and then diluted them to OD_600_ of 0.1 in 2 mL of TSB in 10 mL culture tubes. Two sets of triplicates per strain were prepared, and 5 μg mL^-1^ HQNO was added to one set. A total of 200 μL from each prepared set of triplicates were added to a 96-well plate. To avoid evaporation, the use of the culture wells on the edge of the plate was avoided, and sterile water was added to the empty wells around the experimental wells so that all wells had 200 µL liquid to minimize evaporation. The 96-well plate and lid were sealed with parafilm around the meeting joint to avoid evaporation. The strains were grown in a Biotek Epoch 2 Microplate Reader for approximately 16 hours with constant shaking (shake speed 2) at 37°C, and OD_600_ readings were taken every 15 minutes. The culture optical densities (A_600_) were standardized to the media blanks, and data were plotted as log10 culture optical density vs. time. Two data points were picked that lay within the linear region (logarithmic growth) that were taken one hour apart (t1 and t2), and growth rates were determined using the following equation:

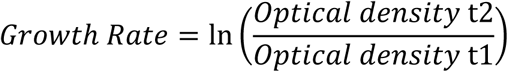

### Transcriptional reporter assays

Strains were grown overnight in TSB containing 10 µg mL^-1^ Cm and subsequently diluted to a 0.1 (A_600_) in 2 mL of TSB-Cm in 10 mL culture tubes. After 8 hours of growth, 200 μL of each sample were added to black 96-well plates (Thermo-Scientific), and fluorescence emission was measured at a 488 nm excitation, 510 nm emission, using a Varioskan Lux plate reader (Thermo Scientific). For assays with *P. aeruginosa* spent medium, *P. aeruginosa* strains were grown overnight in 3 mL of LB in a 30 mL culture tube. Cultures were centrifuged at 13,000×g for 1 minute, followed by filter sterilization of supernatants using 0.22 μM syringe filters (VWR). After *S. aureus* dilution, 100 μL of the *P. aeruginosa* spent media was added to 2 mL *S. aureus* culture. For assays utilizing HQNO, a final concentration of 0-10 μg mL^-1^ (typically 5 μg mL^-1^) of HQNO was added to *S. aureus* cultures that had been diluted to 0.1 (A_600_) in 2 mL of TSB-Cm in 10 mL culture tubes. Sample fluorescence was standardized to the culture optical densities (A_600_) at harvest time.

### Assessment of membrane potential

Overnight cultures were diluted 1:100 in a 10 mL culture tube with 2.5 mL of TSB and grown for 6 hours at 37 °C degrees with constant shaking. Two mL of cells were pelleted by centrifugation. Cells were washed twice with PBS before diluting cells to an optical density (A_600_) of 0.085, and 200 μL were added in triplicate to black 96-well plates. 3,3’-Diethyloxacarbocyanine Iodide (DiOC_2_(3)) was added to a final concentration of 1 mM. The plate was incubated statically at 37 °C in the dark for 30 min. Fluorescence was then measured (450 nm excitation, 650 nm emission).

### Oxygen consumption rate

To assess oxygen consumption rate (OCR) samples were analyzed using a Seahorse XFe96 Extracellular Flux Analyzer (Agilent Technologies) as previously described (52). Briefly, Overnight cultures were diluted 1:100 into 10 mL of TSB media in 125 mL flasks and cultured with and without 5 μg mL^-1^ HQNO at 37°C with shaking at 300 rpm for 4 hrs. Cells were then diluted to a 0.025 OD_600_ and 200 μL were added into each well of a 96-well cell culture microtiter plate. The Seahorse analyzer was calibrated by hydrating the XFe sensor cartridge with sterile water overnight in a non-CO_2_ incubator at 37 °C. Pre-warmed XFe calibrant was added to the wells 2 hours before measurement. The OCR and ECAR were measured through 15 cycles with 3 minutes of measurements and 3 minutes of mixing.

### Metabolomics and sample preparation

#### Sample preparation

Triplicates of bacterial strains were cultured overnight in TSB before diluting to 0.1 (A_600_) in 5 mL of TSB in 25 mL culture tubes. Cells were grown for an additional 8 hours, and 1 mL of each sample was centrifuged at 13,000×g for 1 minute and washed once with PBS. Cell pellets were resuspended in 1 mL of an ice-cold 2:2:1 solution of Methanol:Acetonitrile:H_2_O. Harvested cells were lysed in a FastPrep homogenizer (MP Biomedicals) with 600 µL of 0.1 mm lysing matrix beads (MP Biomedicals) (2 cycles, 40 sec, 6.0 m sec^-1^). Samples were incubated in ice for 5 minutes before centrifuging twice at 13,000×g for 2 minutes at 4 °C. A total of 850 µL of the supernatant was added to a spin filter and centrifuged again at 4 °C 13,000×g for 15 minutes. The samples were frozen at -80°C until they were transferred to the metabolomics core of the Rutgers Cancer Institute of New Jersey for analysis.

#### UHPLC chromatography conditions

The HILIC separation was performed on a Vanquish Horizon UHPLC system (Thermo Fisher Scientific, Waltham, MA) with XBridge BEH Amide column (150 mm × 2.1 mm, 2.5 μm particle size, Waters, Milford, MA) using a gradient of solvent A (95%:5% H_2_O:acetonitrile with 20 mM acetic acid, 40 mM ammonium hydroxide, pH 9.4), and solvent B (20%:80% H_2_O:acetonitrile with 20 mM acetic acid, 40 mM ammonium hydroxide, pH 9.4). The gradient was 0 min, 100% B; 3 min, 100% B; 3.2 min, 90% B; 6.2 min, 90% B; 6.5 min, 80% B; 10.5 min, 80% B; 10.7 min, 70% B; 13.5 min, 70% B; 13.7 min, 45% B; 16 min, 45% B; 16.5 min, 100% B and 22 min, 100% B (53). The flow rate was 300 μl min^-1^. Injection volume was 5 μL and column temperature was 25 °C. The autosampler temperature was set to 4°C and the injection volume was 5µL.

#### Full scan mass spectrometry

The full scan mass spectrometry analysis was performed on a Thermo Q Exactive PLUS with a HESI source which was set to a spray voltage of -2.7kV under negative mode and 3.5kV under positive mode. The sheath, auxiliary, and sweep gas flow rates of 40, 10, and 2 (arbitrary unit) respectively. The capillary temperature was set to 300°C and aux gas heater was 360°C. The S-lens RF level was 45. The m/z scan range was set to 72 to 1000m/z under both positive and negative ionization mode. The AGC target was set to 3e6, and the maximum IT was 200ms. The resolution was set to 70,000.

#### Data analysis and quality

The full scan data was processed with a targeted data pipeline using MAVEN software package (54). The compound identification was assessed using accurate mass and retention time match to the metabolite standards from the in-house library.

Prior to running the samples, the LCMS system was evaluated for performance readiness by running a commercially available standard mixture and an in-house standard mixture to assess the mass accuracy, signal intensities and the retention time consistency. All known metabolites in the mixture are detected within 5ppm mass accuracy. Method blank samples matching the composition of the extraction solvent are used in every sample batch to assess background signals and ensure there isn’t carryover from one run to the next. In addition, the sample queue was randomized with respect to sample treatment to eliminate the potential for batch effects.

### S. aureus survival assays

The survival assays were prepared by growing overnight cultures of *S. aureus* in 2 mL of TSB in 10 mL culture tubes and *P. aeruginosa* strains in 3 mL of LB in a 30 mL culture tubes. These cultures were then diluted 1:100 in triplicate in their respective growth media and cultured for an additional 2.5 hours. Culture optical densities (A_600_) were then standardized to an optical density (A_600_) of 0.4 in one mL of TSB. The *P. aeruginosa* strains were further diluted 1:1000 (1 µL) into the culture tubes containing one mL of a *S. aureus* strain (OD 0.4). The bacteria were then co-cultured at 37 °C for 24 hours with agitation. After 24 hours, the co-cultures were centrifuged at max speed for two minutes, and the supernatant was discarded. The cell pellets were resuspended in PBS and repeated two additional times. The washed cells’ optical densities (A_600_) were measured and then standardized to an optical density of 0.1 in PBS. A serial dilution up to 10^-7^ was performed for each of the co-cultures, and 5 µL of each dilution was spotted on TSB plates containing 7.5% NaCl to inhibit *P. aeruginosa* growth. The plates were incubated at 37 °C for 24 hours, and colony-forming units (CFU) were determined.

### Quantitative real-time PCR assay

Bacterial strains were grown overnight in TSB and diluted to an optical density (A_600_) of 0.1 into 5 mL of TSB with or without 5 μg mL^-1^ HQNO in 30 mL culture tubes in triplicate. Cells were grown for 8 hours at 37 °C with constant agitation, after which they were pelleted by centrifugation 13,000×g for 1 minute. Cell pellets were treated with RNAprotect (Qiagen), and washed in 0.5 mL PBS, pH 7.4, and resuspended in 100 μL of 50 mM Tris, pH 8, containing 6.7 μg lysostaphin and 6.7 μg DNAse. Cells were then incubated at 37 °C for 30 minutes. A total of 200 μL of a lysis buffer solution (20 mM sodium acetate, 1 mM EDTA, 0.5% SDS, 13.4 μg lysostaphin) was added to the samples and incubated at 65 °C for 5 min. RNA extraction and isolation were done as previously described (55). The cDNA library was constructed using the High-Capacity cDNA Reverse Transcription Kit (Biosystems). A StepOnePlus thermal cycler (Applied Biosystems) performed the quantitative real-time PCR. The transcripts were standardized to transcripts corresponding to the 16S RNA gene (SAUSA300_1841; *rrsC*). All datasets were analyzed using the comparative C_T_ method (39, 56).

## Acknowledgments

This project was supported by NIAID 1R01AI139100-01 and USDA MRF project NE−2248 to JMB.

## Table Legend

**Table S1. Abundances of metabolites associated with the WT after culture with and without 5 μg per mL of 2-Heptyl-4-Quinolone N-Oxide (HQNO).**

